# Fossilized Microbial Forms in Baltic and Goan Amber-A Comparative Pioneer Study

**DOI:** 10.1101/209973

**Authors:** Dabolkar Sujata, Kamat Nandkumar

## Abstract

This paper is based on surveys, exploration and standardization of techniques to recover rare amber samples from sands in Goa and identify specimens embedded with distinct microbial fossils based on studies on reference samples of imported Baltic amber. We developed techniques to locate, detect and identify amber samples in local sand. In this pioneer study, we report presumptive microbial forms such as actinobacteria and fungi in amber samples of Goa throwing light on microbial paleobiodiversity. Baltic amber (Succinate) is fossilized resin belonging to the Eocene period (44-49 million years old) derived from the Araucariaceae and Leguminoseae families of trees containing up to 8% of Succinic acid and compounds such as terpenoids and phenolic derivatives. Sooty moulds in the amber fossils have been studied (Schmidt et al., 2014). Samples of imported Baltic amber were validated, studied and used as reference for identification and characterization of amber found in sand of Goa. FTIR Spectroscopic tests diagnostic of presence of Succinate proved that both samples meet the criteria as plant derived Succinate containing products. Fossil fungi in Baltic amber were observed, and compared with similar forms in rare amber fragments of Goa. These samples were subjected to microscopic examination. Actinobacterial and fungal forms embedded in local amber were compared with similar forms found in imported Baltic amber and those published in literature. Detection of fossilized actinobacterial and fungal forms has shown us the potential for further studies for comprehensive collection and microscopic examination of such paleomicrobial forms in rare local amber samples.

## INTRODUCTION

Amber (Succinate=C_6_H_6_O_4_) is the fossilized resin produced from the trunks and the roots of certain trees mainly belonging to family Pinaceae. Plant that secreted Baltic amber has been already identified as *Pinites succinifer.* Amber is a complex mixtures of diverse compounds, such as terpenoids and phenolic derivatives (Poinar and George, 1992) is found distributed globally in Russia, France, Germany, Lebanon, Spain, Dominican Republic, Austria, USA, Myanmar, Japan, and Mexico (Schmidt et al., 2014). The age of all amber specimens has been determined to be minimum 4 to maximum 300 Million years old. Amber primarily yellow in colour displays shades ranging from light yellow to dark yellow to brown to black (Aranda et al., 2007). Nathaniel Sendel (1686 - 1757) is pioneer in research on plants, microbes and animals embedded in Baltic amber. Plants secret resin when they suffer injury. The biota such as microbes, plant parts and even animals gets trapped inside this resin.

Over the years, the process of fossilization occurs and resin is converted into amber and the bio inclusions remains inside the amber (Pontin et al.,2000). Organisms that were trapped in resin, and are preserved in amber, are called bioinclusions and some of these are especially informative about various taphonomic processes, paleoenvironmental conditions and important paleobiological aspects (Speranza et al., 2015). Amber has been used since prehistory in the manufacture of jewelry and ornaments. Scent of amber and amber perfumery is well known. Amber oil has been use for medicinal purpose (Poinar and George, 1992). India subcontinent separated from Gondawana land around 100 mya and collided with Asia 52 millions years ago (Veevers et al., 1995). The formation of Indian amber took place probably during Eocene period ie 53 million years ago (Rust et al., 2010). Fossils records show that Coniferopsida such as *Buriadia, Walkomiella, Searsolia and Paranocladus* resin producing trees were once part of lower Gondwana (Pant, 1982). These resin deposits could be present as amber in placer deposits and beach sands of Goa which was part of Gondwana. Recent advances in spectrometry allow the development of physical characterization of amber to confirm that amber originates from various kinds of plants such as conifers and *Fabaceae* (Langenheim, 2003). Differences between “true/natural/biogenic” and false/synthetic/artificial amber can be identified by some classical tests for example true amber produces sweet, pine smell when burnt and does not dissolve in acetone (Pionar, 1992). True amber floats in salt water and this called as salt water test (Raducanu, 2006). Amber burns with a black smoke like incense and does not melt and this forms the basis of validation of amber by burning test (Raducanu, 2006). Other tests of amber analysis include florescent test, refractive index test, IR spectroscopy, polarized light test (Raducanu, 2006; Ascaso etal., 2007). Various scientists (Stubblefield et al., 1988; Pionar & George, 1992; Pontin et al., 2000; Smejkal et al., 2011; Hartl et al., 2015) have reported presence of paleomicroflora in the amber, particularly interesting finds are of sooty mould morpho species (Schmidt et al., 2014).

**The main objectives of this study were**

I. Studies on validated Baltic amber samples imported in Goa as standard reference material
II. Specific studies for detection of bioinclusions in Baltic amber samples
III. Development of simple techniques to harvest and identify amber from local sand samples
IV. To compare the bioinclusions in Baltic and local amber with special stress on fossilized microbial morphotypes
V. To gain insights into paleomicroflora (eubacteria, actinobacteria, fungi) of Goa

## MATERIALS AND METHODS

### Baltic amber

The majority of Baltic amber from Russia is dated to Eocene period (44-49 million years old). Areas of Baltic amber deposition and Baltic sea is shown in the figure 1. The Baltic amber samples used for this study were officially imported from Russia in Goa for sale to tourists. Higher flow of tourists (32392 in 2006 to 162746 in 2013 and 104890 in 2014–2015) from Russia and Ukraine to Goa by chartered flights has also introduced many traded commodities like semi precious stones and Baltic amber. Since amber of this quality was not easily available in India the Baltic amber traded in Goa was utilized for scientific studies. Identification of specimens with bioinclusions was carried out, further confirmation and validation was done by FTIR spectroscopy. Microscopic studies of the amber samples were carried out using light microscopy, Phase contrast microscopy and photomicrography.

**Fig 1:**
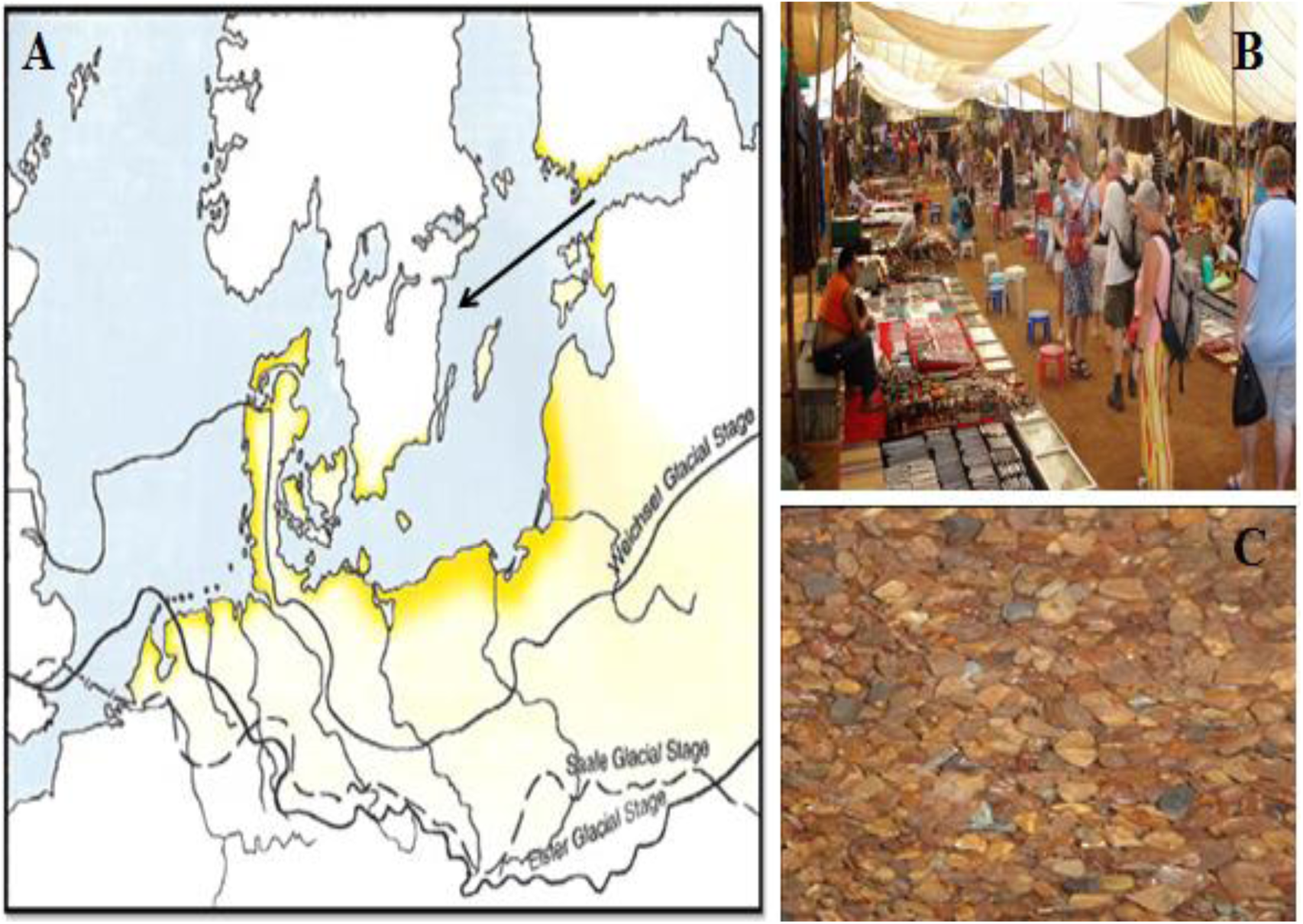
A-Areas of Baltic amber deposition and Baltic sea (Poinar & George, 1992); B-Baltic amber samples imported from Russia and marketed in local markets; C-Baltic amber collection.

### Indian amber

Goa is the second smallest state located on the west coast of India covering an area of 3702 sq kms and runs 105 km long and 65 km wide. Goa is located between latitudes 14º 53’ 55’’N and 15º 47’ 59’’N and longitudes 73º 40’ 34’’E and 74º 17’ 03’’ E. The Arabian Sea marks the western boundary of the State. Selection of sampling sites was carried out using Google earth (Fig 2). Sands samples were collected from placer deposits and intertidal zones of beaches by pool sampling methods. Separation and sieving of sand samples were carried out in the laboratory. In a second step, using stereomicroscope, hand-picked amber specimens were separated, were confirmed and validated by acetone test (Poinar & George, 1992). Stereoscopic screening of one kg (about 500 plus) of specimens were carried out. These specimens were further confirmed and validated by FTIR spectroscopy using standard FTIR spectra for Baltic amber as reference. Microscopic studies of the amber samples were carried out as mentioned above. Identification of specimens with bioinclusions was carried out using standard pictorial keys on paleobioforms reported in amber (Poinar & George 1992; Schmidt et al., 2014).

**Fig 2:**
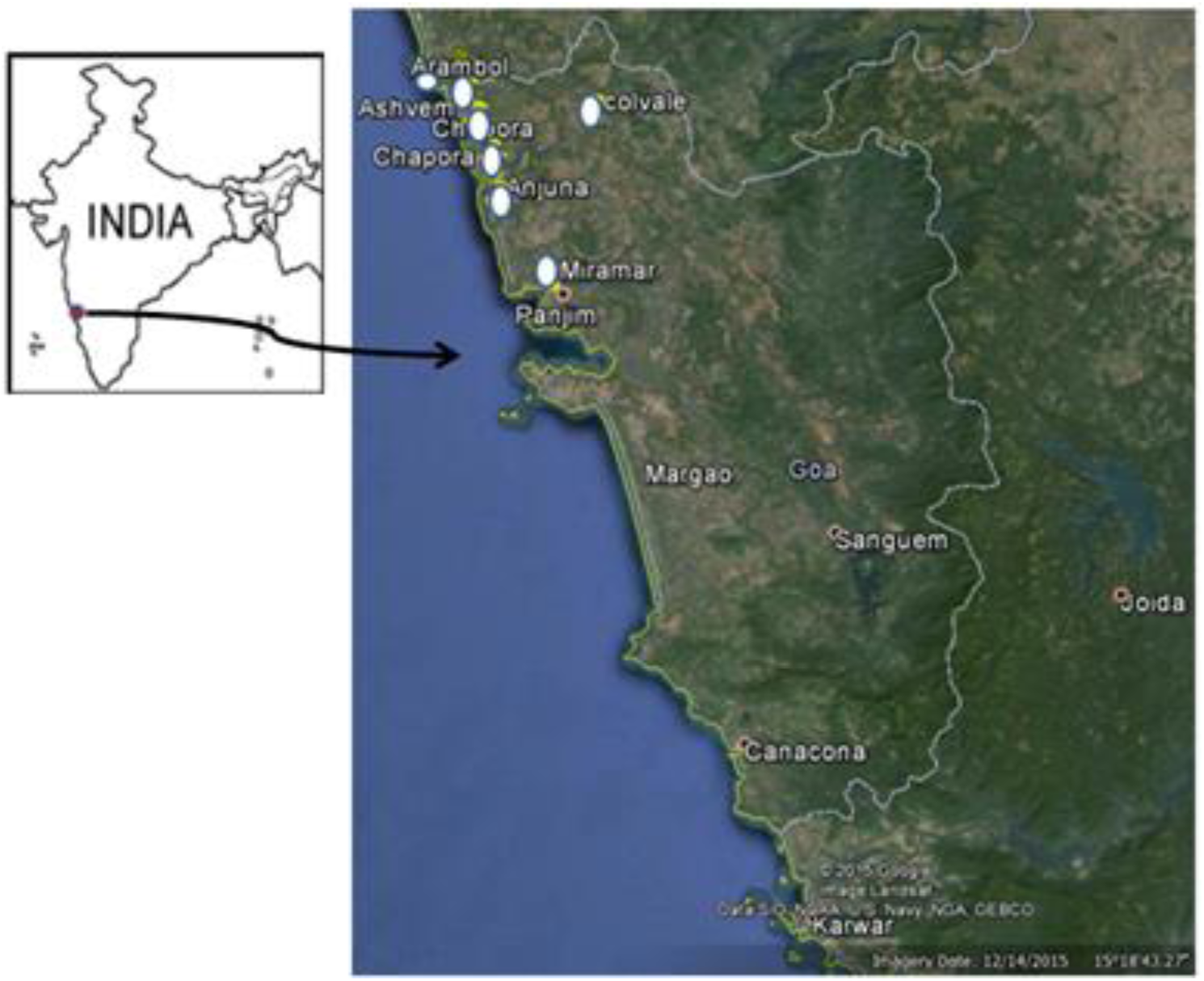
The Google earth image locating the sampling sites. White circle indicates the location of beaches of Goa.

**Fig 3:**
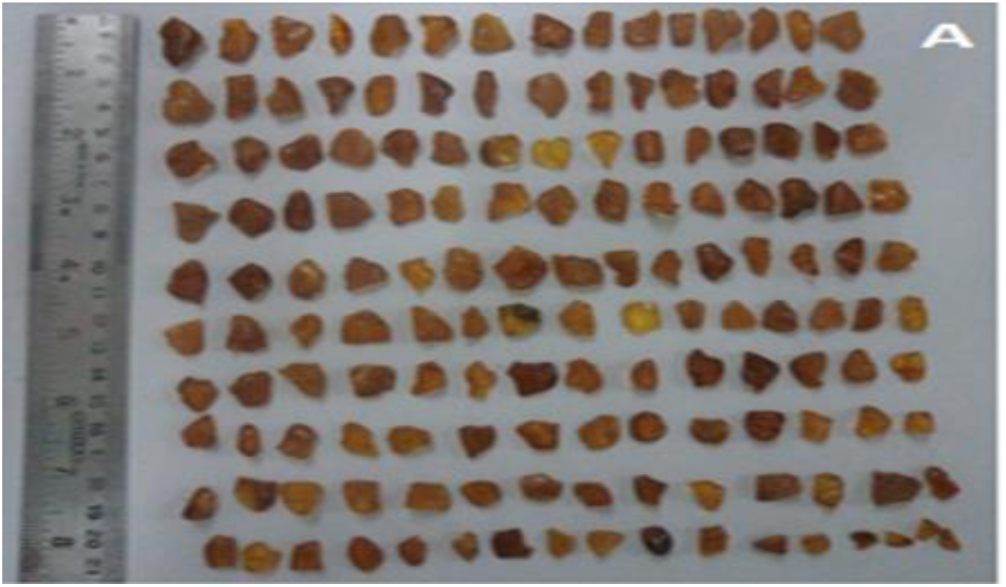
Baltic amber specimens.

## RESULTS

### Microscopic studies of Baltic amber

The size of the Baltic amber specimens ranged from 1-2cms. Baltic amber showed the presence of bacterial and fungal bioinclusions. Bacterial inclusions included spores, actinobacterial substrate hyphae, and thin filaments. Fungal inclusions included hyphae, codidiophores and conidia, sporangia, and septate fungal spores. Coccoid bacterial spores having the diameter of 0.5–0.8 μm (fig 4A) were found dispered in amber. A few amber specimens showed the presence of thick actinobacterial spiral hyphae and a number of long, thick fungal hyphae (fig 4B, C). Baltic amber also showed presence of characteristic thick microfungal conidiophores and sporangia(Fig 4D) and entrapment of septate fungal spore chains (Fig 4E).

**Fig 4(A-D):**
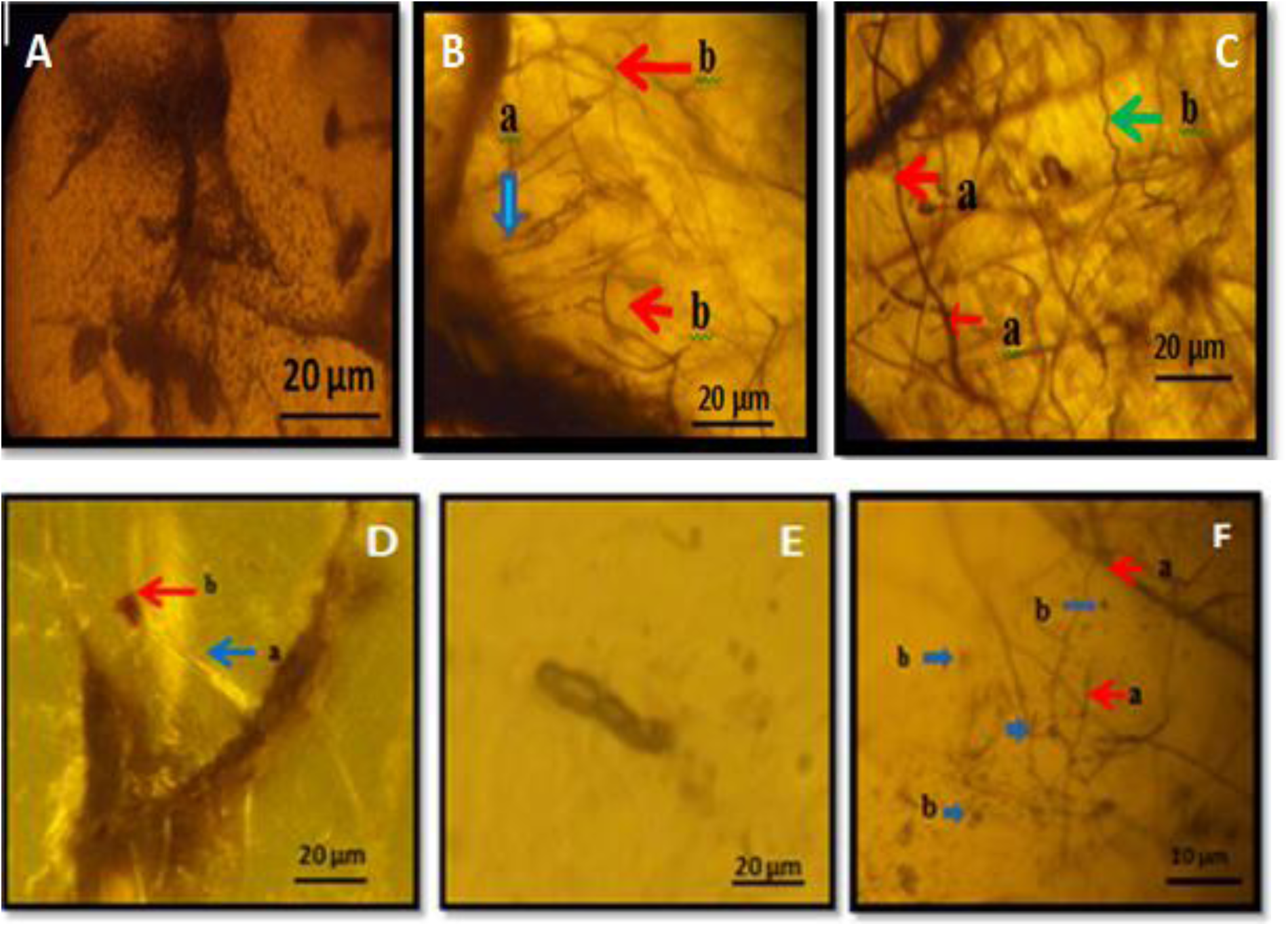
A-Coccoid spores dispersed in amber; B-Actinobacteria (Blue arrow)–a, spiral hyphae, b-fungal hyphae (Red arrow); C-Fungal Hyphae (Red arrow-a) and Actinobacterial filaments (Green arrow-b); D-microfungal Conidiophore (Blue arrow-a) and sporangia (Red arrow-b) of microfungi, E-Septate fungal spore chain, F-Micro fungal Hyphae (Red arrow-a) and Micro fungal spores (Blue arrow-b).

### Microscopic studies of Indian amber

The size of the amber from Indian sands ie from beaches of Goa ranged from 40-70 μm. The colour of these amber specimens ranged from light yellow to dark orange. The sieving procedure helped to reveal the interesting microscopic fragments of amber surviving in local sand samples (Fig 5). As indicated in fig 6 local amber fragments showed the presence of wide inflated thick fungal hyphae, basidiomycetes microcolony, dark mitosporic fungi with bulbous hyphae all of which remain unidentified.

**Fig 5(A-H):.**
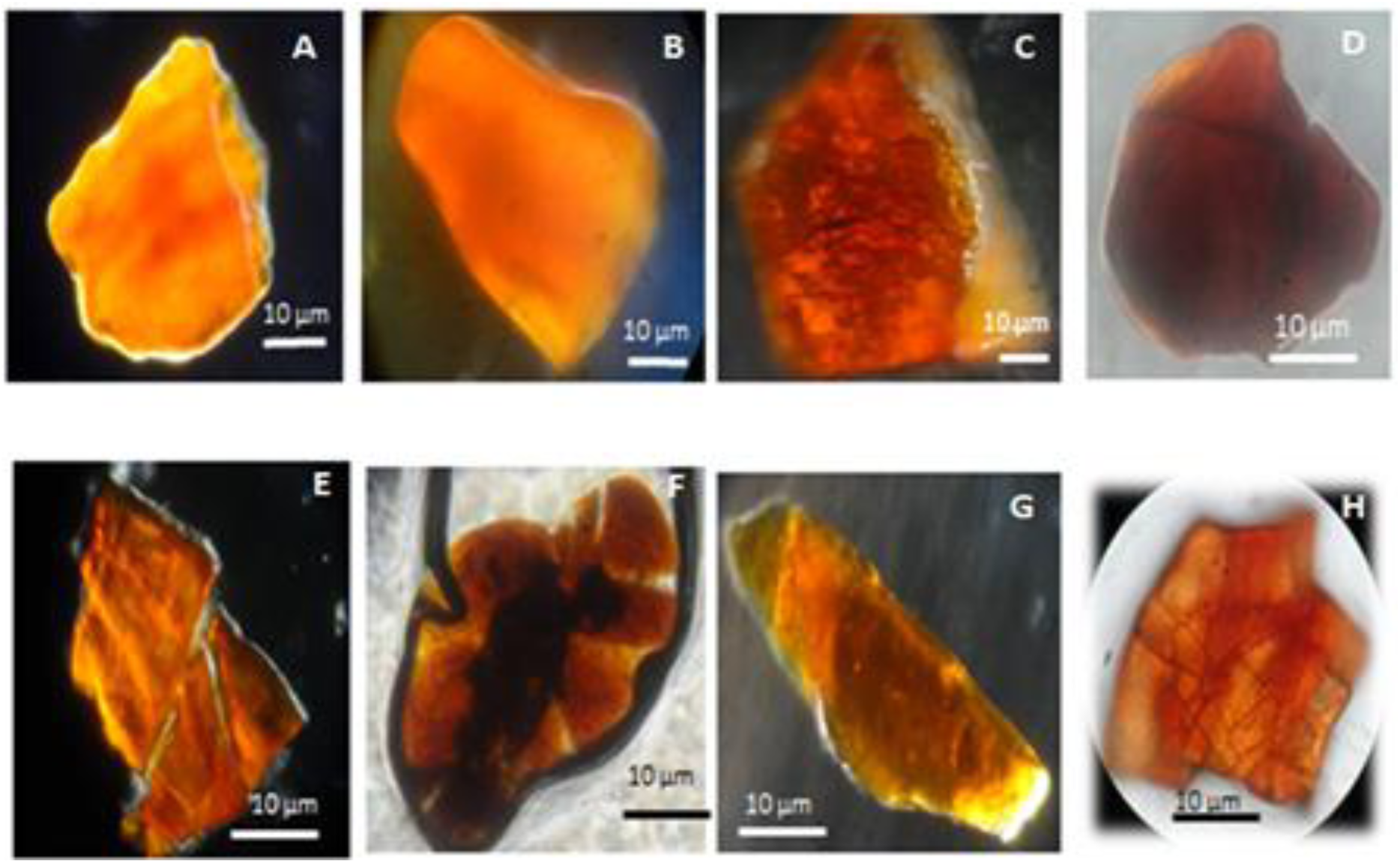
Interesting microscopic fragments of amber surviving in local sand samples

**Fig 6(A-F):**
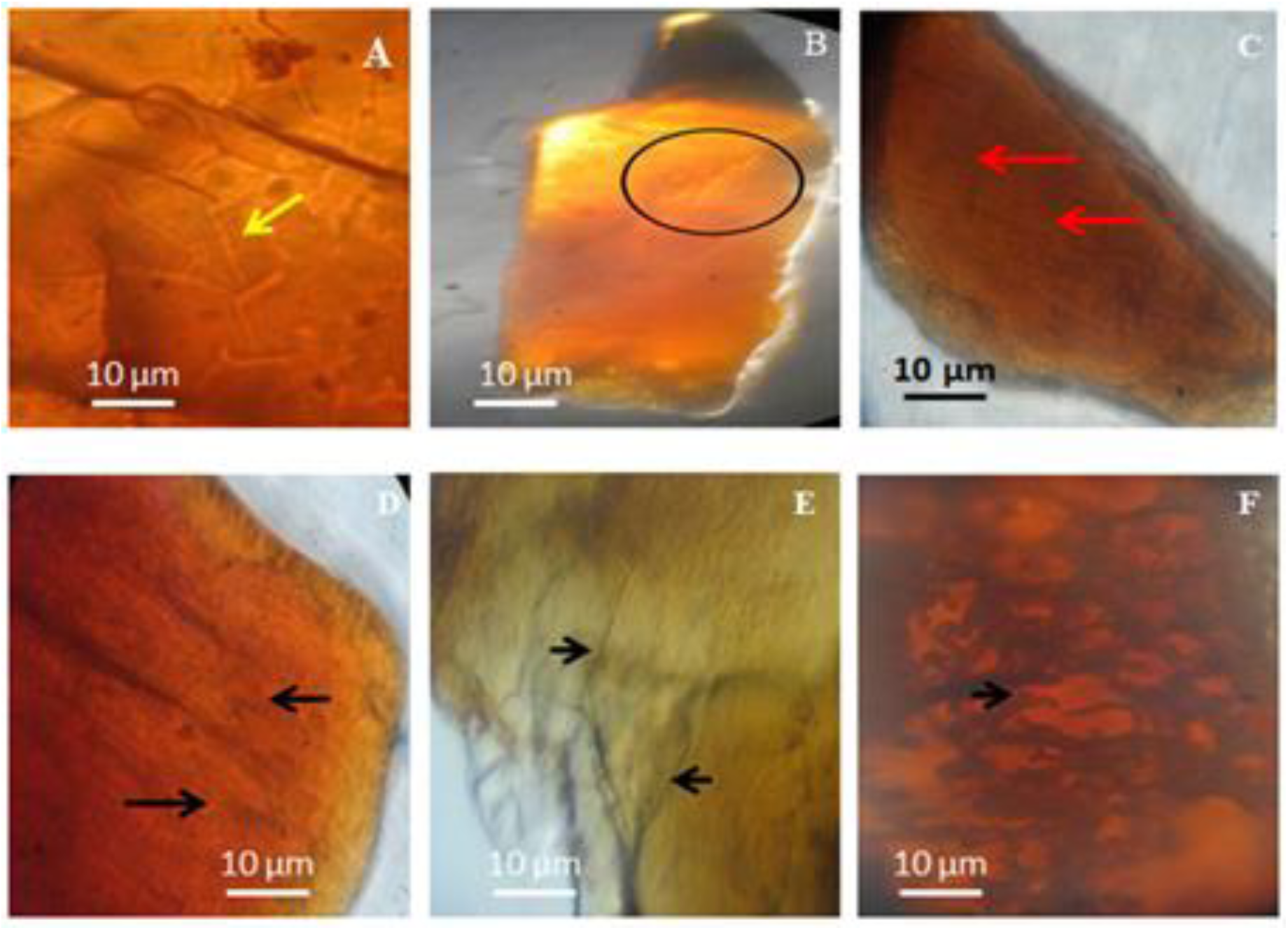
Bioinclusion in Goan microamber. A-Wide thick fungal hyphae; B-Basidiomycetes colony; C-Basidiomycetes colony; D- Basidiomycetes colony; E- Basidiomycetes colony; F-Dark mitosporic fungi with bulbous hyphae.

## DISCUSSION

This is the first study of imported Baltic amber in India and claimed to be the first modest study to report detection of bioinclusions. This is the first report on recovery of micro amber or fragments of amber from coastal sands of Goa, west coast and south India. Earlier studies on Indian were on bacterial filament from an enigmatic piece of amber from Assam-Arakan Basin of eastern India (Girard et al., 2014) and well preserved in amber-embedded biota such as fungal bodies has been reported from lower Eocene(~52 Ma) formation of Gujarat (Mukherjee et al., 2005). Baltic and Indian amber showed difference in sizes. Baltic amber specimens were fairly microscopic ranged from 1-2 cms whereas Indian amber under light microscope ranged from 40-70 μm indicating their relative rarity. We designate these forms as ‘microamber’ or disintegrated forms of amber (DFA). Conditions under which Baltic amber may be found are different than conditions in tropical India. Genesis of microamber in sand may be specific to coastal erosive tropical environment. Disintegration of original large pieces of amber may take place because of demanding hot, humid, turbulent and erosive tropical conditions in India. Large pieces of amber can be easily disintegrated due to natural forces such as abrasion, friction, erosion, action of turbulent currents (Poinar & George, 1992). In natural sandy strata abarasion due to shifting sand particles which have sharp edges may also cause disintegration over period of time. The sand sieving procedure helped in enriching tiny fragments of amber. Out of all the processed fragments or microamber 70% showed presence of unidentified microbial inclusions where as other 20% showed unidentified plant and animal inclusions and 10% were void with no inclusions. Simillar observations were noticed by Mukherjee et al., 2005 in amber from lower Eocene in Gujarat. Both Baltic and Indian amber specimens showed the presence of bacterial and fungal bioinclusions. However their origin may be different (Pionar &George, 1992; Rust et al., 2010). The microbial bioinclusions in amber in Goa could shed light on paleobiodiversity of plant community and associated microflora.

## CONCLUSIONS

Detection of fossilized actinobacterial and fungal forms has shown that there is potential for further studies for comprehensive collection and microscopic examination of such entombed microbial forms in rare local microamber samples. *Micrococcus luteus* and *Bacillus sphaericus* has been successfully isolated from million years old Dominian amber (Cano, 1995 and Greenblatt et al., 2004). Similar attempts to isolate pure cultures aseptically from local microamber specimens are in progress. If these succeed then molecular and genetic studies of these pure isolates will be carried out. Bioprosepcting of these cultures may provide important biotechnological leads. Systematic isolation of microamber from different locations and recovery of useful ancient micro-organisms from these may open a new and exciting area of amber microbiology in India. It has not escaped our notice that metagenomic approach may also be possible if DNA from the microamber in Goa could be recovered by using bioinclusions as positive proxies for presence of genetic material (Cano, 1995; Greenblatt et al., 1999; Lambert et al., 1998; Smejkal et al., 2011).

## ACKNOWLEDGEMENTS

This work was supported by UGC-SAP Phase II–Biodiversity, Bioprospecting programme and forms part of ongoing work of Goa University Fungus Culture Collection and research unit (GUFCCRU) On mapping microbial biodiversity. We thank R.N.S Bandekar CO, Vasco Da Gama for partial funding.

